# Development of Kinetic Modeling to Assess Multi-functional Vascular Response to Low Dose Radiation in Leukemia

**DOI:** 10.1101/633644

**Authors:** Jamison Brooks, Bijender Kumar, Darren M. Zuro, Jonathon D. Raybuck, Srideshikan Sargur Madabushi, Paresh Vishwasrao, Liliana Echavarria Parra, Marcin Kortylewski, Brian Armstrong, Susanta K Hui

## Abstract

Vascular permeability, tissue transfer rate (K_trans_), fractional extracellular tissue space (ν_ec_) and blood perfusion are crucial parameters to assess bone marrow vasculature (BMV) function. However, quantitative measurements of these parameters in a mouse model are difficult because of limited resolution of standard macroscopic imaging modalities. Using intravital multiphoton microscopy (MPM), live imaging of dextran transfer from BMV to calvarium tissue of mice bearing acute lymphoblastic leukemia (ALL) was performed to obtain BMV parameters. Mice bearing ALL had increased BMV permeability, altered K_trans_, increased ν_ec_, decreased blood perfusion, and increased BMV permeability resulting in reduced drug uptake. Targeted 2 Gy radiation therapy (RT) to mice bearing ALL increased local BMV perfusion and ALL chemotherapy uptake (P<0.0001 and P=0.0036, respectively), suggesting RT prior to chemotherapy treatment may increase treatment efficacy. Developed MPM techniques allow for a quantitative assessment of BMV functional parameters not previously performed with microscopic or macroscopic imaging.

## Introduction

Bone marrow is a highly vascularized organ, capable of rapid vascular changes to support bone growth, maintenance, and hematopoiesis [1]. Bone marrow vasculature (BMV) is essential for the delivery of oxygen and nutrients to both bone and bone marrow. Alterations in BMV function have been shown in hematological malignancies, tumor bone metastasis, and bone diseases by macroscopic time-lapsed imaging techniques such as DCE-MRI and DCE-CT [2–5]. These imaging based compartmental modeling parameters, such as the kinetic transfer constant (K_trans_) and fractional extracellular extravascular tissue space (ν_ec_) have predictive value for both the diagnosis and prognosis of bone marrow related disease. K_trans_ and ν_ec_ are influenced by the functional characteristics of individual vessels such as permeability, density, morphology, and single vessel blood perfusion. Malignancy-related vessel remodeling can result in a variety of changes to these vessels including angiogenesis, increased vessel permeability, and vasoconstriction [6, 7]. As a result, many malignant environments are both hypoxic and have poor drug delivery, making both chemotherapy and radiation therapy treatments less effective [8–10]. Vascular function has been shown to be important in slowing the onset of disease and assisting with the treatment of hematological malignancies [7, 11], suggesting the importance of BMV in the growth and treatment of malignancies. A functional characterization of BMV is therefore crucial to understand the healthy bone marrow environment, as well as the functional changes that occur with the onset of disease and treatment.

Macroscopic blood perfusion imaging techniques assess the overall efficiency of tracer delivery to the tissue of interest. However, macroscopic imaging modalities, such as magnetic resonance imaging, X-ray based computed tomography, or whole animal fluorescence imaging do not have the resolution required to identify features from single capillary-sized blood vessels, making observations of the underlying functional vascular changes difficult or impossible [12]. The resolution limitations also cause difficulties in modeling tracer uptake in the bone marrow of a mouse model, further limiting usefulness in a preclinical setting. Alternatively, most microscopic imaging modalities are able to identify the functional changes in individual vessels, but have not, to our knowledge, been used with kinetic compartmental modeling to obtain quantitative parameters such as K_trans_ and ν_ec_, which allow for better understanding of vascular dynamics with the onset of disease.

To address the need for a single imaging platform capable of BMV kinetic modeling at the necessary resolution to distinguish capillary sized vessels, we developed multiphoton microscopy (MPM)-based time lapsed imaging of BMV in the calvarium of live anesthetized mice. Time lapsed images of fluorescent dextran leakage from the vascular lumen to the interstitial tissue were fit to a compartmental kinetic model to quantitatively assess K_trans_, ν_ec_, and vessel permeability (P) while also monitoring single vessel blood perfusion, functional vessel surface area, vascular morphology, and vascular density at cellular resolution. Mice bearing GFP+ Philadelphia chromosome B-cell acute lymphoblastic leukemia (ALL) were chosen as a model, since ALL-induced vascular alterations have been shown through various signaling pathways [13, 14]. Additionally, little work has been done showing ALL-induced functional alterations of BMV with live in-situ imaging. We administered targeted local image guided radiation therapy (IGRT) to ALL-bearing and healthy mice to modulate the local BMV and validate compartmental modeling-based fitting parameters, respectively. Hoechst and daunorubicin uptake on IGRT treated and untreated regions of mice bearing ALL were assessed with the hypothesis that low dose (2-4 Gy), locally targeted IGRT treatments to regions of leukemic burden can enhance local drug uptake through the alteration of BMV and reduction of leukemic burden.

## Results

### BM microenvironment remodeling and leukemia localization during ALL growth

MPM imaging revealed leukemic homing out of BMV to the bone marrow within 30 minutes of ALL injection (Supplemental Video S1). ALL cells were observed flowing rapidly through BMV, as well as rolling on and adhering to the endothelial wall. Progression of ALL growth was tracked by the acquisition of large tiled three-dimensional z-stack images (**Error! Reference source not found**.A-F, Video 1). Degradation of the frontal bone suture near the bregma was present in mice at high ALL burden (Figure 1G-J) suggesting leukemia-induced bone reduction and tissue destruction. In the case of degraded calvarium bone, the majority of ALL cells remained under the bone-collagen rather than moving into the expanded suture space. X-ray computed tomography (CT) scans revealed a reduction in the Hounsfield Units on the frontal lobe of ALL mice as compared to healthy control mice, likely because of a decrease in bone mineral density (Figure 1K).ALL was primarily observed in regions between vasculature and bone-collagen and was predominately homogenously distributed throughout calvarium marrow space (Figure 1L). However, occasional regions of the calvarium did contain heterogeneous pockets of ALL (**Error! Reference source not found**.M). No obvious changes in ALL distribution were observed from mice imaged 2 to 13 days post ALL injection except for an overall increase in the density of ALL (**Error! Reference source not found**.N).

**Figure 1:**
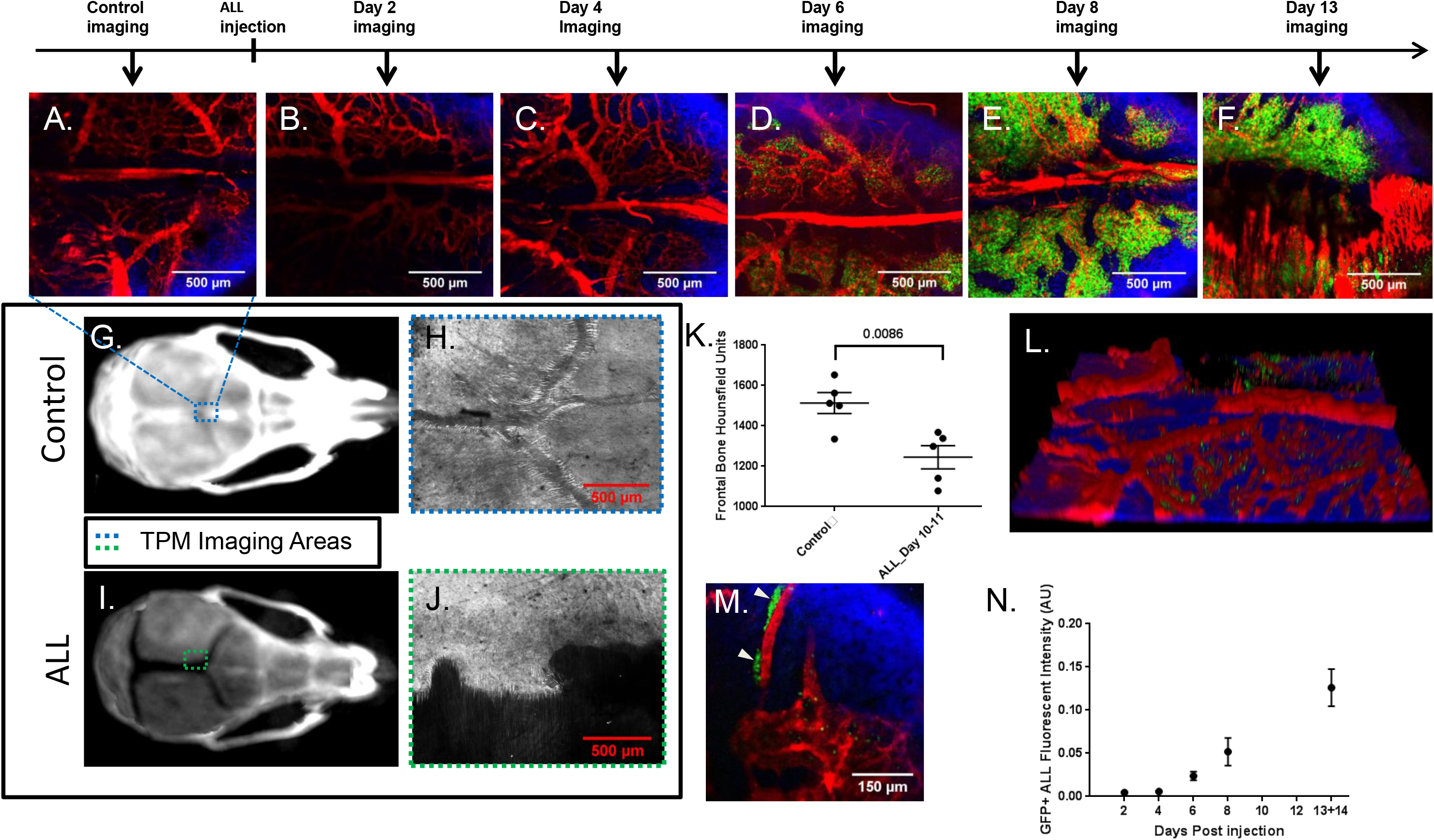
MPM tile imaging of ALL progression and bone degradation in mouse calvarium. Tiled MPM images of the calvarium frontal bone in healthy control mouse (A) and images from 2, 4, 6, 8, and 13 days post BCR-ABL injection are shown (B-F). Red is vasculature, blue is bone-collagen, and green is GFP ALL signal. CT images of control and ALL bearing mice 12 days post ALL injection (G, I). Dotted lines indicate relative regions of MPM imaging. Tiled MPM images of bone-collagen second harmonic generation are shown from separate control and ALL bearing mice 14 days post injection (H, J). CT Hounsfield Units from contoured frontal bones of healthy control and ALL bearing mice 10-11 days post injection of ALL (K). A 3D volume rendering of a tiled mouse skull 4 days after injection of ALL showing the vasculature and ALL on the underside of the bone-collagen is shown (L). Heterogeneous pockets of ALL in calvarium located in close proximity to both BMV and bone-collagen are marked with white triangles (M). GFP+ ALL fluorescent intensity for individual mice and their respective days post injection (n=4, 4, 4, 3, 3 mice for each time point respectively) (N).

### Non-linear changes in K_trans_ and increased BMV permeability with ALL burden

The first pass of dextran post injection was observed by MPM time-lapsed imaging as early as 6 seconds post injection. Dextran had reached the capillary bed within 12 seconds for both healthy and ALL bearing mice. Dextran blood plasma concentration was notably increased on the first pass before complete diffusion into the blood (Video 2). In all cases, dextran blood plasma concentration was well described by bi-exponential decay (Supplemental Figure 1A-B), suggesting consistent perfusion to BMV from the arterial blood pool. Dextran uptake into calvarium bone marrow tissue started immediately after entrance into the blood (Figure 2A-H & Supplemental Figure 1C-D). Dextran tissue kinetics showed good agreement with the compartmental model used, and no obvious cellular uptake of dextran was observed (Supplemental Figure 1E). The two methods for calculation of the transfer coefficient (K_trans_ and K_slope_) showed strong positive correlation with each other (R^2^=0.9059, Supplemental Figure 2A). Validation of the fitting methodology was performed by perturbing the system using targeted image guided radiation therapy (IGRT) in healthy mice, as radiation has been shown to both increase extracellular fraction (because of cellularity decrease), and damage vasculature, causing increased leakage [15, 16]. Four-Gy radiation treatments to the left half of the calvarium and left femur significantly increased ν_ec_ and K_trans_ in IGRT treated-calvarium over that of untreated mice (P=0.0038 and P=0.0305, respectively, Figure 2I-J). A corresponding reduction of cellularity in treated femur and calvarium regions of mice was found as compared to non-treated regions (P=0.0076, ratio paired t-test P=0.02077, Supplemental Figure 2B-C). Measurements of dextran fluorescence of crushed femur supernatants revealed significantly higher fluorescence in IGRT-treated femur supernatants (2.23 ± 0.10×10^6^) compared to untreated controls (0.90 ±0.04×10^6^, P=0.0003, Supplemental Figure 2D), suggesting radiation significantly increases total drug delivery to the bone marrow tissue. A negative correlation between femur cellularity and dextran fluorescence was found, confirming that extracellular space and the total dextran uptake by bone marrow cells are influenced by cellular density (R^2^=0.7735, Supplemental Figure 2E).

**Figure 2:**
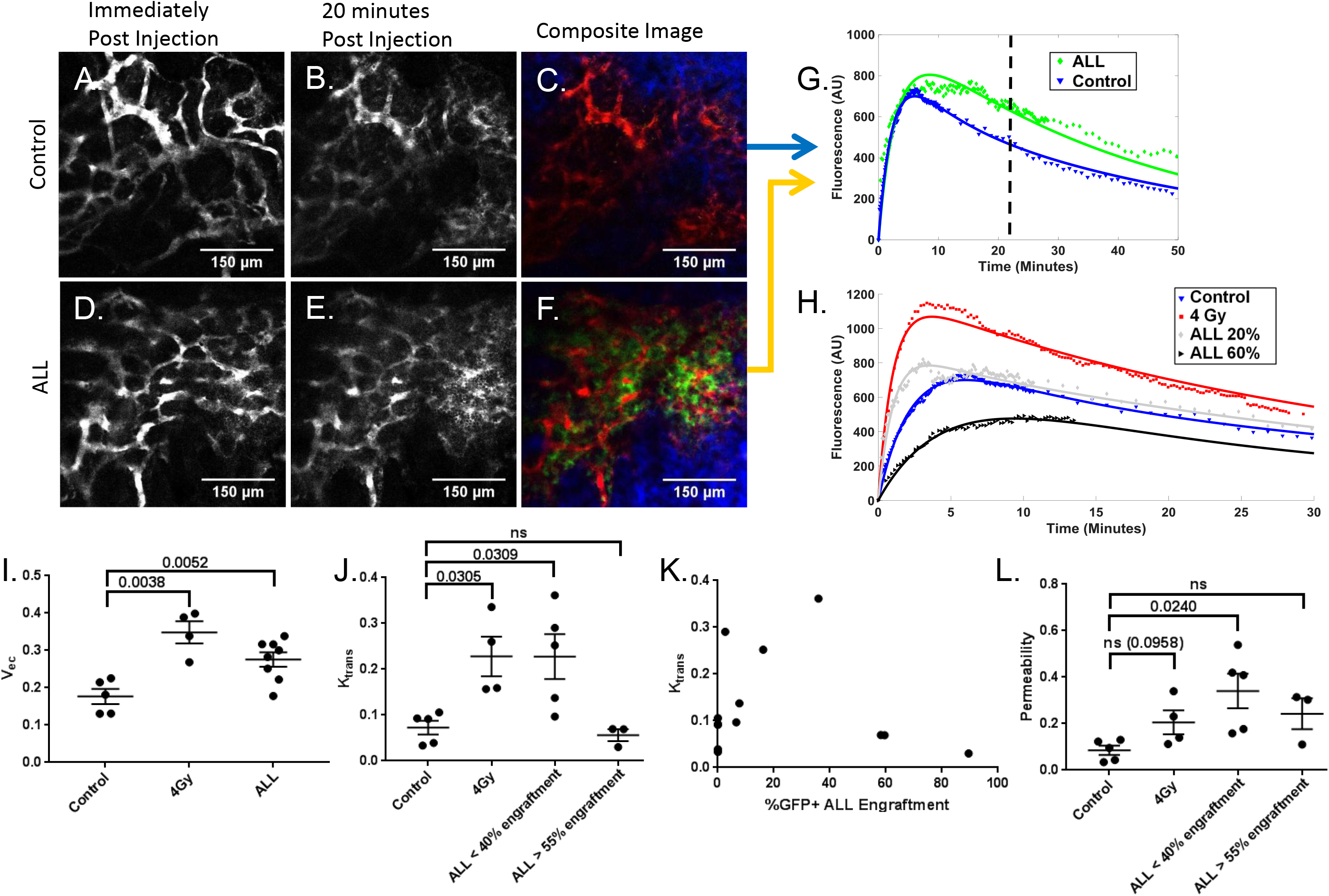
Time lapsed MPM imaging and kinetic compartmental modeling observe changes in K_trans_ ν_ec_, and vascular permeability. Representative images of dextran perfusion imaging are shown for control and ALL mice. Dextran fluorescence is shown for healthy and control mice immediately post injection and twenty minutes post injection (A,B,D,E). Merged images verifying the local presence of ALL(C,F). Green is GFP+ ALL fluorescence, red is dextran fluorescence and blue is collagen bone second harmonic generation. Dextran time lapsed data and fitted tissue compartment curves from matching displayed images are plotted for single healthy control and ALL mice (G). The vertical striped line indicates the 20 minutes post injection depicted in images. Increased tissue uptake in the plotted data and images are seen at 20 minutes in the ALL mouse compared to the control mouse. Dextran tissue kinetics from a single healthy control, non-leukemic 4 Gy treated, low burdened ALL, and high burdened ALL mice are plotted (H). Values for ν_ec_ are shown for healthy control, non leukemic irradiated, and ALL mice (I). Values for K_trans_ and vascular permeability are shown for healthy control, non leukemic irradiated, low engraftment ALL, and high engraftment ALL mice (J, L). A plot of ALL bone marrow engraftment versus K_trans_ for untreated mice is shown (K).

A higher ν_ec_ was found in ALL-bearing mice (0.28±0.02) over that of healthy animals (0.18 ±0.02, P=0.0052, Figure 2I), whereas no significant difference in cellularity was noted (P=0.7524), suggesting that change in ν_ec_ is due to other factors, such as bone degradation and high interstitial fluid pressure. At <40% engraftment, K_trans_ (0.23±0.05) was higher than that of healthy control mice (0.073±0.015, P=0.0309) and decreased back to control levels when leukemic burden exceeded 50-60% (Figure 2J-K). Measurements of BMV surface area per volume (S) from time lapsed images showed a negative correlation between leukemic burden and functional vessel surface area (R^2^=0.7893, Supplemental Figure 2F). A significant increase in vascular permeability was observed in mice with <40% ALL engraftment (0.34±0.07) over that of healthy controls (0.085±0.020, P=0.0240, Figure 2L & Supplemental Figure 2G).

Time lapsed imaging of daunorubicin, a chemotherapeutic agent, was performed in an ALL-bearing mouse and compared to dextran kinetics to ensure that MPM-based imaging can be used for fluorescent agents of varying sizes and pharmacokinetic uptake patterns. A clear distinction between vascular, tissue, and cellular compartments could be visualized with daunorubicin (Supplemental Figure 3 A-J & Supplemental Video S2). A faster peak uptake of daunorubicin (30 s) is observed compared to that of dextran (185 s) because of its smaller molecular weight (Supplemental Figure 3K).

### BMV remodeling results in angiogenesis, poor perfusion, and loss of functional vessels with ALL

MPM-based measurements of calvarium mean vessel density were higher in ALL mice at both 8 days (424±22 mm^−2^) and 12-14 days (606±43 mm^−2^) post ALL injection compared to that of healthy control mice (287±24 mm^−2^, P=0.0061 and 0.0017, respectively, Figure 3A-E). Platelet endothelial cell adhesion molecule (CD31) was used as a marker for endothelial cells, as it is commonly expressed in endothelial cells and involved in new vessel formation [17]. Flow cytometry revealed an increase in the CD31+ endothelial fraction of total live bone cells with the onset of ALL over that of control (P=0.0032, Figure 3F & Supplemental Figure4A). No significant differences in total bone marrow cellularity between healthy and ALL bearing mice were observed, suggesting an overall increase in the total number of endothelial cells present in mice bearing ALL (P=0.7524, Supplemental Figure 4B). Flow cytometry measured a significant reduction in high endomucin-expressing endothelial (CD31+) cells in mice 13-14 days post ALL injection (P<0.0001, Figure 3G Supplemental Figure 4C). These cells have previously been reported to co-localize with osteoblast progenitors in the growth plate regions of the femoral bone and is responsible for hematopoietic stem cell (HSC) maintenance by secreting HSC-supporting cytokines [18]. Data suggest that ALL transforms the hematopoietic niche into an environment more favorable for ALL at the expense of normal hematopoietic cells by altering the endothelial subset. Similar alterations have been seen in acute myeloid leukemia [7, 11]. Endothelial expansion was accompanied by large changes in vascular function and vascular morphology. MPM revealed a significant reduction (8.93 μm) in average vessel inner diameter of mice 12 -14 days post ALL injection compared to control (P<0.0001, Figure 3H). Average velocities of cells flowing through the BMV network of ALL bearing mice (1.34±0.11 mm/s) were reduced compared to healthy control mice (3.70±0.484 mm/s, P<0.0001). Healthy untreated control mice typically had small inner diameter vessels with rapid blood flow connected to large inner diameter, slow flowing vessels. Alternatively, mice with moderate to high ALL burden displayed a more homogenous population of vessels in terms of both blood flow velocity and vessel diameter, suggesting that typical cellular extravasation, oxygen delivery, and drug delivery may be altered in mice bearing ALL because of a loss of vasculature phenotype (Figure 4A-H).

**Figure 3:**
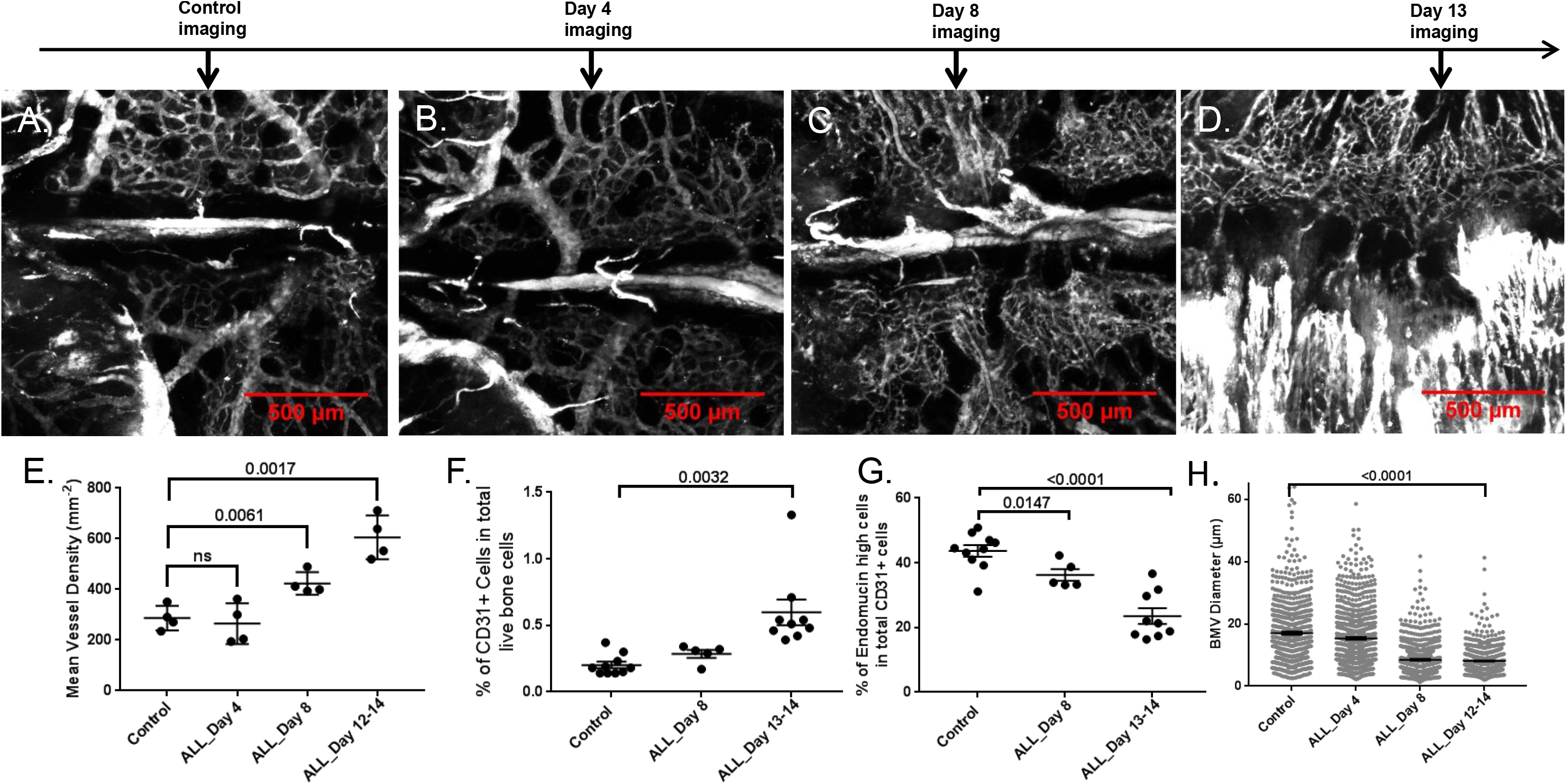
BMV angiogenesis and remodeling occur with the onset of ALL. Representative maximum intensity projection tiled MPM images of vasculature for control (A), 4 days (B), 8 days (C), and 13 days post ALL injection(D) are shown. Angiogenesis and morphology changes can be observed. Mean vessel density measurements of healthy control mice and BCR-ABL mice 4, 8, and 12-14 days post ALL injection (E). Bone marrow flow cytometry analysis of the percentage of CD31+ endothelial cells in total live bone cells for control, 8 days, and 13-14 days post ALL injection (F). The fraction of high endomucin expressing endothelial cells in total endothelial cells in bone marrow for control, 8 days post injection, and 13 days post injection is shown (G). Measurements of mean vessel diameter for control and ALL burdened mice 4, 8, and 12-14 days post injection are shown(n=3-4) (H).

**Figure 4:**
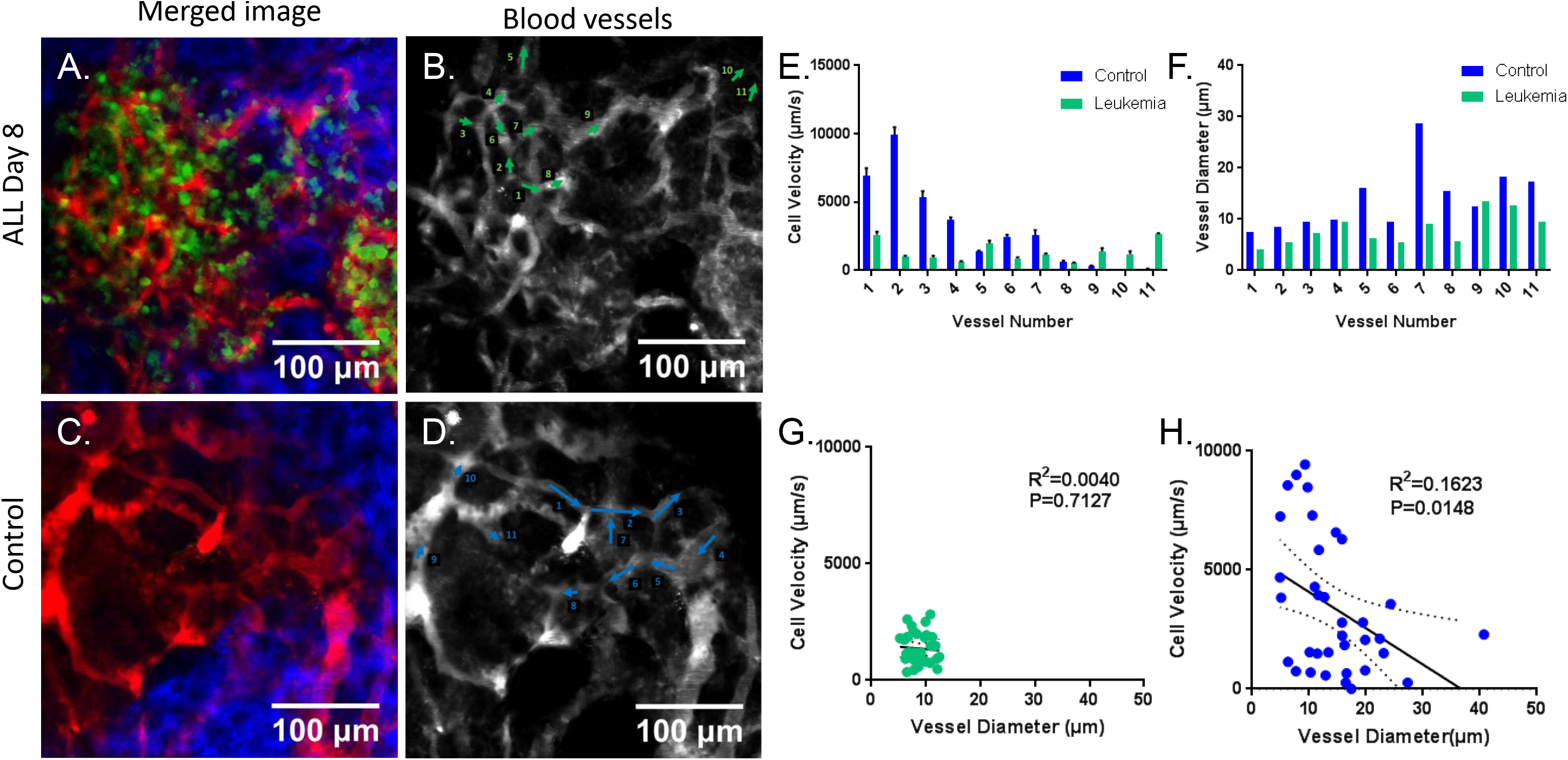
ALL induced changes to BMV alter the vascular structure and reduce blood perfusion rates. Merged color images of ALL burdened (A) and control mice (C). Green is GFP+ ALL signal, red is vascular lumen fluorescence and blue is collagen bone second harmonic generation. Vascular blood pool images with corresponding blood vessel mapping of upstream to downstream vessels labeled 1-11 for ALL burdened vessels 8 days post ALL injection (B) and healthy control vessels(D). The matching vessel flow (E) and diameter (F) measurements from each individual vessel show the loss of upstream fast flow small diameter vessels and large diameter downstream slow flow vessels in ALL. Plots of cell velocity versus diameter for ALL burdened vessels eight days post transplant (G) and healthy control vessels (H) along with their non-zero significance and R-squared values are shown. Individual vessel measurements from 3 separate mice were used to acquire for both control and ALL datasets.

Pockets of semi-collapsed vessels were observed in mice as early as day 8 post ALL injection (Supplemental Figure 5A-B). Accumulation of stalled non-moving cells in blood vessels was commonly observed near regions of partial vascular collapse (Supplemental Figure 5C). Time-lapsed imaging of fluorescent dextran showed BMV opening and closing intermittently in mice bearing ALL (Video 3). This phenomenon was not observed in healthy control mice (Video 4). In mice with high leukemia burden, vessels closed off after 15-20 minutes, preventing longer time lapsed permeability measurements (Supplemental Video S3). Unidirectional movement of GFP+ ALL toward regions of vascular collapse was observed in time lapsed images prior to vessel collapse, suggesting extravascular fluid flow influences cellular position and vessel function. Measurements of systolic and diastolic pressure were taken to better understand the mechanics behind vascular closing. Reductions in systolic (56.2 mm Hg) and diastolic pressure (31.0 mm Hg) were observed with the onset of leukemia (P=0.0002 and 0.0375, respectively, Supplemental Figure 6A-D), suggesting that reduced intravascular blood pressure may also be playing a role in vessel collapse.

### Restoration of perfusion and chemotherapy delivery using local, anatomically targeted IGRT in ALL

Local IGRT treatments to half of the calvarium and a single femur altered vascular morphology and function of BMV in mice bearing ALL (Figure 5A-E). Two-Gy local IGRT treatments to BMV of mice bearing ALL significantly increased vascular diameter (13.4 μm) and the average velocity of cells flowing through the BMV network (2.41 mm/s) compared to values in untreated mice bearing ALL (P< 0.0001 and 0.0001, respectively Figure 5F-H). Two-Gy local IGRT treatments restored average cellular vascular velocity to a level similar to that in healthy untreated control mice (P=0.9474,). A significant decrease in average total bone marrow cellularity (~14 million cells) and average bone marrow ALL burden (15.4%) was found with 4 Gy treatments (P=0.0029 and 0.0196, respectively Figure 5I-K). Hoechst dye and daunorubicin chemotherapy were used to test drug delivery with IGRT-treated and untreated regions of mice bearing ALL. The percentage of total cells stained positive for Hoechst in untreated calvarium regions was shown to have a strong negative correlation (R^2^=0.965) with ALL engraftment (Figure 6A), while no significant changes in overall cellularity were seen in the bone marrow (P=0.7524), suggesting that cellular uptake is inhibited by the onset of leukemia. Flow cytometry revealed an increase in the percentage of ALL cells labeled positive for both Hoechst (19.7%) and daunorubicin (18.5%) in 2 Gy IGRT treated regions compared to that in untreated regions (P=0.0121 and 0.0036, respectively, Figure 6B-F & Supplemental Figure 7A-B). Similar trends were observed in 4 IGRT treated ALL bearing regions. Healthy control mice showed high percentages of Hoechst positive cells in total cells for both 2 Gy IGRT treated (96.1±1.2%) and untreated calvarium (92.3±5.3%, Supplemental Figure 7C). MPM imaging of mice bearing ALL observed revealed a higher daunorubicin fluorescence in 2 Gy IGRT treated calvarium than in untreated calvarium (Figure 6G). ALL Cellular segmentation showed significantly increased cellular uptake of daunorubicin in the 2 Gy-treated calvarium (0.038 ±0.002) region compared to the untreated region (0.031±0.002), consistent with flow cytometry data (P=0.0362, Figure 6H-I). Mice bearing ALL treated with 2 Gy IGRT showed somewhat higher percentage of ALL cells labeled positive for Hoechst (92.2±2.3 %) than mice bearing ALL treated with 4 Gy IGRT (86.8±0.9%, P=0.1242), despite the fact that untreated calvarium regions in 2 Gy treated mice had higher ALL engraftment (32.2±5.5 %) than in untreated calvarium regions in 4 Gy-treated mice (25.8 ± 1.1 %, P=0.3604). Additionally, 2 Gy-IGRT treated bone marrow regions had increased cellularity (~10 million cells) compared to that from 4 Gy IGRT (P=0.0136). Data suggests 2 Gy may be a more optimal dose to condition BMV for chemotherapy.

**Figure 5:**
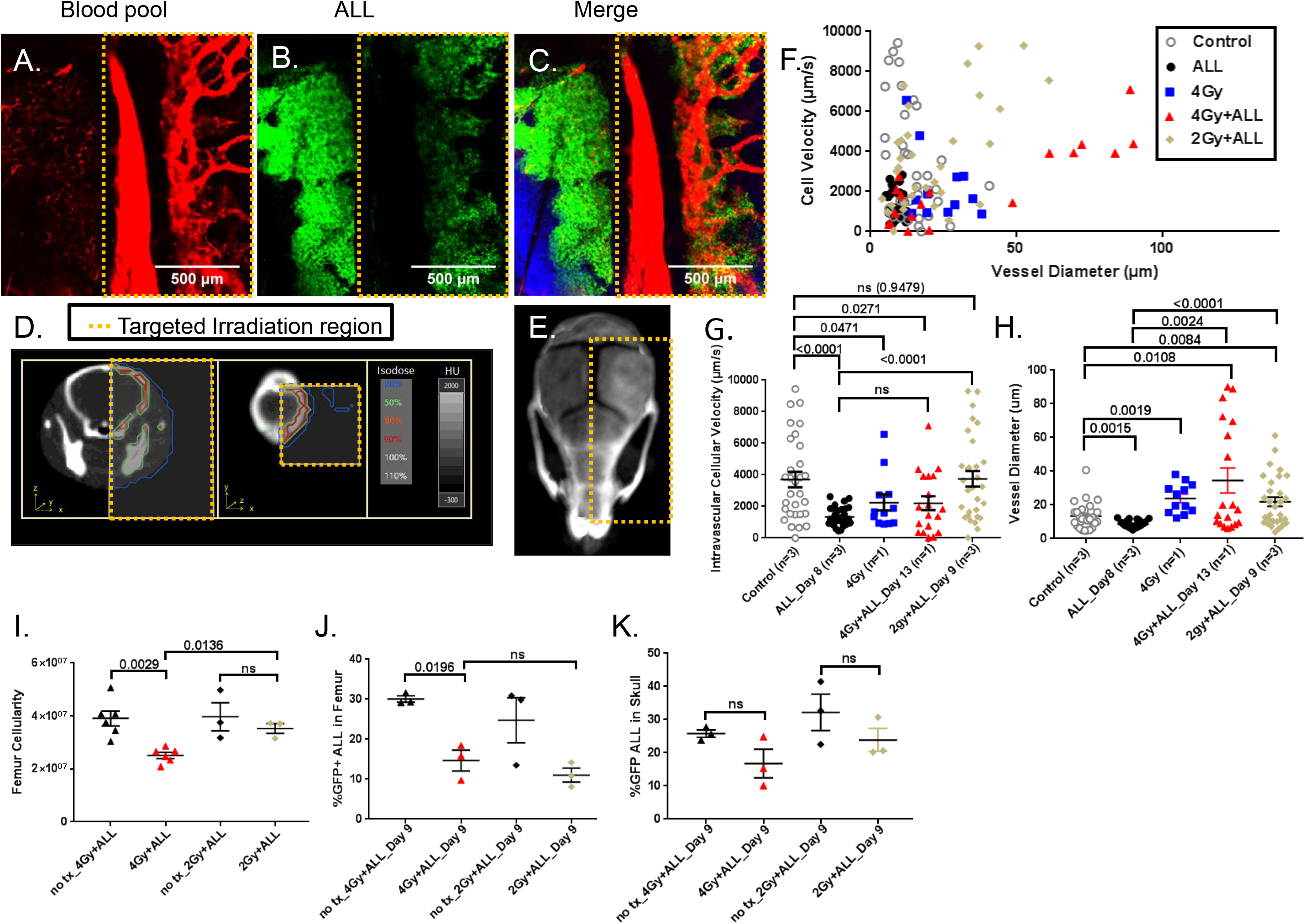
Low to moderate dose IGRT induces vasodilation and restores blood perfusion. Tiled MPM images taken 2 days post 4 Gy IGRT and 13 days post ALL injection, depicting treated and non treated regions of the mouse calvarium blood pool (red), ALL (green), and merged image with colored bone-collagen second harmonic generation (blue) (A-C). CT slice images showing treated regions and their corresponding relative isodose lines are shown (D). A three-dimensional rendered CT based volume image with the targeted treatment region is shown (E). Vessel diameter and cellular velocity plots of individual blood vessels are plotted for healthy control mice (n=3), non treated ALL bearing mice (n=3), 4Gy IGRT treated healthy mice (n=1), 4Gy IGRT treated ALL bearing mice(n=1), and 2Gy IGRT treated ALL bearing mice (n=3). 4Gy and 2Gy IGRT treated ALL mice were imaged 13 days and 9 days post ALL injection respectively. There are 30, 30, 12, 19 & 30 individual vessel measurements for each group respectively (F-H). Femur total cellularity, femur ALL engraftment and skull ALL engraftment of treated and non treated regions for 4Gy IGRT treated healthy control, 4Gy IGRT treated ALL, and 2Gy IGRT treated ALL mice 9 days post ALL injection (I-K). All IGRT treatments were performed 40-48 hours prior to imaging or tissue harvest.

**Figure 6:**
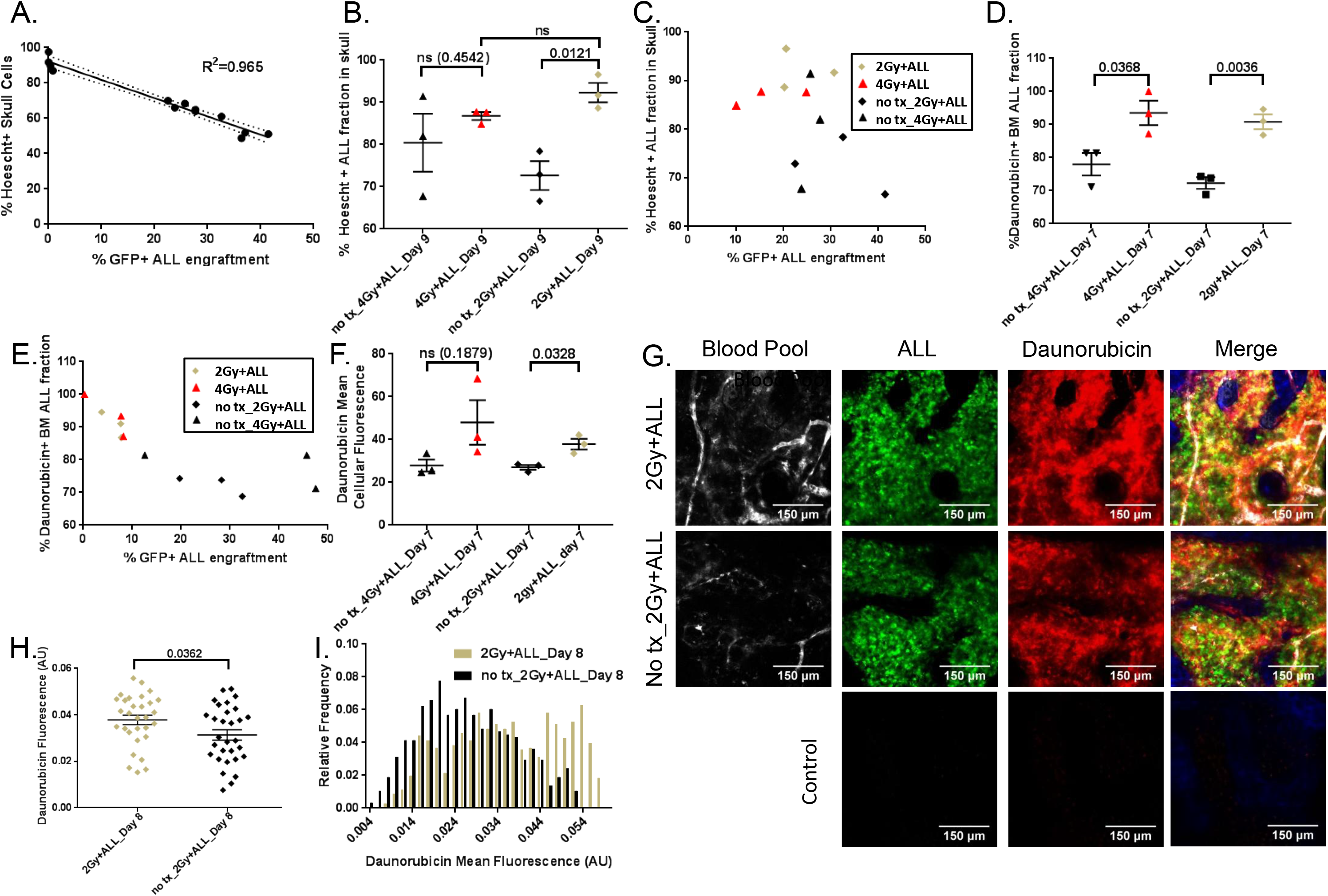
Low to moderate dose IGRT enhances ALL cellular uptake of Hoechst and daunorubicin. Flow cytometry analysis of the percentage of total cells stained positive for Hoechst in untreated skull are plotted with GFP ALL engraftment (A). Flow cytometry analysis of the percentage of ALL cells stained positive for Hoechst and daunorubicin are plotted for 4Gy IGRT treated regions, 2Gy IGRT treated regions, and corresponding non treated regions (B-E). Flow cytometry based mean cellular daunorubicin fluorescence of ALL cells for 4Gy IGRT regions, 2Gy IGRT regions, and corresponding non-treated regions (F). Three mice were used for each matching treated and untreated regions. MPM Images of treated and non treated regions of a mouse receiving 2Gy IGRT to half of the calvarium and images of a healthy control mouse receiving no injections of daunorubicin (G.). Plotted fluorescence of daunorubicin uptake based on image segmentation of single ALL cells for 30 cells with the highest GFP mean fluorescence for shown treated and non treated images (H). Relative frequency histogram plots of all ALL segmented cells from both treated and non treated images (I).

## Discussion

In this study, quantitative compartmental modeling was developed for MPM time lapsed imaging to measure increased BMV permeability, the nonlinear response of K_trans_, increased ν_ec_, reduced vascular perfusion, and altered vascular morphology in mice bearing ALL. ALL-induced alterations resulted in reduced cellular drug uptake. Targeted low-dose IGRT treatments to regions of leukemic burden were given to ALL-bearing mice with the hypothesis that treatments would alter BMV and enhance local drug uptake. Low dose IGRT increased vascular perfusion and enhanced chemotherapy delivery to mice bearing ALL, suggesting that targeted low dose irradiation prior to standard induction chemotherapy may result in increased treatment efficacy against ALL and potentially other hematologic malignancies.

Increased vascular permeability results in decreased BMV functional efficiency, adversely impacting drug delivery in mice bearing ALL. At low to moderate leukemic burden, K_trans_ is increased compared to healthy mice, likely because of increased vascular permeability, enabling greater dextran transport through the endothelial wall. However, at high leukemic burden, vascular remodeling results in collapsed vessels, reduced perfusion, and reduced vascular surface area, which lowers K_trans_ back to values similar to those of control mice. Increased K_trans_ values in low burden ALL may indicate a therapeutic window of enhanced delivery for ALL antibody-based therapies such as those targeting CD19 or CD22 which are similar in size to 150 KDa dextran [19, 20]. However, differences in vascular transport mechanisms of dextran and therapeutic antibodies make further assessments of antibody delivery kinetics necessary [21]. Previous reports have shown arterial and sinusoidal blood vessels to be linked with phenotypes observed in healthy mice. Small diameter, fast flow BMV correlates with arterial vessels, whereas large diameter, slow flow BMV correlates with sinusoidal and venule vessels for healthy mice [22, 23]. The reduction in intravascular cellular flow velocity and BMV diameter with the onset of ALL results in a loss of the healthy BMV physiology. Increased interstitial fluid pressure is due in part to increased vessel permeability in many solid tumor models [24, 25] and is likely present in ALL because of increased BMV permeability. Observations of extravascular cellular movement toward BMV just prior to vessel collapse suggest that increased vascular permeability results in increased interstitial fluid flow. Interstitial fluid flow contributes to the intermittent opening and closing of collapsed vessels by moving cells in the calvarium toward or away from regions of BMV. The increased shear force of extravascular fluid flow on accumulated extravascular cells collapses vessels. Accumulation of intravascular cells occurs near semi-collapsed vessels until intravascular pressure increases enough to open vessels, allowing cells in the blood to transiently pass through collapsed BMV. The observed phenomena in conjunction with poor blood perfusion demonstrate the inefficiency of blood transport in the BMV of mice bearing ALL, resulting in poor drug delivery. Early preservation of the healthy vascular system by the inhibition of vascular remodeling *in-vivo* has been shown to give a survival advantage for mice treated with induction chemotherapy, while inhibition of vascular remodeling alone at late stages of disease burden has not shown an effect [11]. The inefficient BMV system also contributes to increased hypoxia, which has been shown to be present in multiple hematological malignancies [10]. Restoration of BMV perfusion and reduction of high disease burden is crucial to improving chemotherapy delivery to the bone marrow.

In contrast to many drug based therapies, radiation treatments have the ability to deliver highly conformal treatments to various regions of high disease burden, while largely sparing the surrounding healthy tissue [26]. Highly conformal low-to moderate-dose radiation treatments in ALL bearing mice alter local vessel blood perfusion and result in a population of dilated, increased flow vessels. These vessels are not seen in healthy untreated, or ALL-bearing untreated mice, and demonstrate the ability of radiation to increase BMV perfusion and dilate collapsed vessels. Similar vascular alterations using radiotherapy (RT) in solid tumors have been reported, potentially because of nitrous oxide dependent vasodilation [27]. RT in combination with therapeutic agents for leukemia that frequently elicit off target tissue toxicity [28, 29] should be explored because of the conformal targeted treatment of RT and its drug delivery enhancing properties. IGRT treatments of BMV suggest that 2 Gy rather than 4 Gy treatments may be better at restoring perfusion and vessel diameter to levels similar to those of healthy untreated control mice. Additionally, Hoechst and daunorubicin delivery to 2 Gy-treated mice bearing ALL was similar to 4 Gy-treated mice bearing ALL, despite the reduced reduction on overall cellularity by 2 Gy treatments. These results suggest further dose optimization should be performed for quantitative correlation between RT timing, dose, and fractionation scheme. MPM measurements will be essential to further optimize BMV perfusion, permeability, and morphology measurement parameters for RT-based combination therapies.

Kinetic compartmental modeling using MPM has the ability to clearly separate the tissue and vascular compartments to obtain K_trans_ and ν_ec_. MPM can also simultaneously reveal the underlying vascular changes that affect K_trans_ and ν_ec_, such as vascular perfusion, vascular morphology, and vascular permeability. Changes in these underlying parameters occur early in the progression of malignant disease, making ν_ec_ and K_trans_ useful as potential biomarkers and for assessing drug or oxygen delivery. Both K_trans_ and ν_ec_ have been clinically used as predictive markers of disease and treatment response, demonstrating their relevance in prognosis and treatment evaluation [2–4, 30].Limited work has been done with microscopy to perform tissue compartment modeling to identify vascular permeability [12, 22]. Moreover, to our knowledge there have been no successful attempts to fit time-lapsed microscopy data of local plasma and tissue concentrations to perform compartment modeling and obtain K_trans_ and ν_ec_ for the local vasculature. Compartmental tissue modeling has only been used in the context of arterial blood to tissue transport with macroscopic imaging modalities [31, 32], which accurately evaluates k_trans_ and ν_ec_ to whole tissues. In contrast, MPM is used to assess tracer transfer from local vasculature to the surrounding tissue while being able to clearly distinguish capillaries from interstitial tissue. The increased resolution of MPM increased resolution allows for direct observation of the underlying vascular changes that influence K_trans_ and ν_ec_, which is often not possible with macroscopic imaging. Additionally, MPM can measure tissue vascular surface area, which allows for the isolation of vascular permeability from K_trans_. The increased resolution of MPM also enables the assessment of vessel parameters near small heterogeneous regions of disease which are present in many hematological diseases [7, 11]. MPM-based compartmental modeling therefore represents a new and robust method for assessing the state of BMV.

## Methods

### Mice, cell lines, and drug treatments

All procedures for animal experimentation were performed according to City of Hope guidelines and approved by the Institutional Animal Care and Use Committee. C57BL/6J mice (Strain 000664, Jackson Laboratory, Bar Harbor, ME) mice were injected with 1×10^6^ GFP^+^ BCR-ABL (p190Kd) expressing B-cell acute lymphoblastic leukemia cells [33] suspended in 200 μl of phosphate buffered saline (PBS) through the tail vein. The age of mice used for experiment sets is shown in the supplemental methods section. Injection concentrations of 10 mg/kg and 40 mg/kg were used for Hoechst 33342 (ThermoFisher Scientific Waltham, MA) and daunorubicin HCL (Selleck Chemicals LLC Houston, TX), respectively, 20 minutes before being euthanized for tissue processing or imaging. All drugs were administered through the tail vein.

### Surgery, multiphoton image acquisition, and image processing

For calvarium imaging, mice were anesthetized initially with 3% isoflurane at a flow rate of 2 L/min, then maintained with 1.5% isoflurane at a flow rate of 1.5 liters per minute. A stereotactic apparatus with a bite bar was utilized to keep the mouse stable during surgery. The skin on top of the calvarium was removed, and a custom titanium head plate [34] with an inner diameter of 8 mm was fixed to the calvarium using Alpha-Dent Polycarboxylate Cement (Dental Technologies, Lincolnwood, IL). The head plate was then inserted into a custom built heated stage maintained at 37°C allowing for easy viewing of the mouse calvarium. For imaging, a Prairie Ultima multiphoton microscope (Bruker Corporation Billica, MA) was used. Fluorescent excitation was performed using a Chameleon Ultra II tunable Ti:Sapphire laser with 140 femtosecond pulses (Coherent, Santa Clara, CA). Image processing was performed using Fiji/ImageJ [35, 36] and Cell Profiler (v.2.2.0) [37] software. Further details of the imaging acquisition, processing, and segmentation can be found in the supplemental methods section.

### Intravascular cellular velocity measurements

Velocities of cells flowing through the BMV network were calculated in a similar fashion as in previous works [22, 23, 38, 39]. See supplemental methods for details.

### Time lapsed imaging and compartmental modeling

Vascular permeability, K_trans_, and ν_ec_ were measured by time lapsed imaging of 150 kDa TRITC dextran uptake (TdB Consultancy, Uppsala Sweden). Mice were intravenously catheterized and injected with 2 μL of Qtracker™ 655 Vascular Label (Invitrogen, Waltham Massachusetts USA) suspended in 48 μL PBS to identify vasculature for imaging. 500 μg of TRITC dextran suspended in 100 μL PBS was injected, and imaging was started simultaneously. A two-compartment model with fixed equal transport rates in and out of the tissue compartment was used [31] (Supplemental Figure 1E). Curve fitting was performed using Matlab (R2018a 9.4.0.81364, MathWorks Natick, MA) by fitting time lapsed imaging data from blood and tissue regions of individual mice to the compartmental data, yielding K_trans_ and ν_ec_. An additional analysis method of the kinetic transfer coefficient (K_slope_) was performed by taking the slope of the tissue concentration curve immediately following dextran injection, while dextran tissue concentration is minimal. Vascular permeability was determined by dividing K_trans_ by the vascular surface area per image volume. Vascular surface area per image volume was obtained by image measurements in Fiji/ImageJ. A detailed description of image acquisition and processing can be found in the Supplemental Methods.

### Blood pressure measurements

Pulse rate, systolic blood pressure and diastolic blood pressure measurements were acquired by tail cuff method on conscious mice. See supplemental methods for details.

### CT image acquisition and targeted irradiation

Mice treatments using targeted dose calculated IGRT were administered to half of the calvarium and to one femur in order to preserve the systemic function and blood delivery of the vascular system to the tissue of interest. Radiation treatments and CT images were administered and acquired with the Precision X-RAD SMART Plus / 225cx small animal image guided irradiation system (Precision X-Ray, North Branford, CT) [40]. Dose calculations were performed by the small animal radiotherapy planning system [40], using a Monte Carlo dose engine [41, 42]. Imaging and tissue harvest were performed 40-48 hours post treatment. See supplemental methods for more details.

### Flow cytometry

Endothelial cell analysis preparations were followed according to Kumar et al. [43]. In brief, the bone cells were blocked with CD16/32 blocking antibody, followed by staining with CD45 (PE/Cy5,clone 30-F11), Ter119 (PE/Cy5,Ter119), CD31 (PE/Cy7,clone 390), endomucin (A660,eBioV.7C7 ebioscience) antibodies (Bio Legend, San Diego) and AmCyan viability dye for 30-45 min at 4°C in dark. Details of sample preparation for staining, drug uptake study flow preparation, and flow cytometry data acquisition were performed as standard and can be found in the supplemental methods.

### Dextran well plate readings

Well plate fluorescence readings of TRITC dextran in crushed femur supernatants can be found in the supplemental methods.

### Statistical analysis

Unless otherwise noted, significance testing between groups was performed using Welch’s two sided t test with a P value of 0.05 for significance. For linear correlation testing, R squared coefficient of determination tests were utilized. All distribution error bars are displayed as the mean plus or minus one standard error of the mean.

## Acknowledgments

Research reported in this publication is supported by the National Cancer Institute of the National Institutes of Health under award number P30CA033572 and partly supported by National Institutes of Health grant 1R01CA154491-01. The authors wish to thank Michael Farrar (University of Minnesota), the University of Minnesota Imaging Core and James Sanchez for providing cell lines, technique development and manuscript editing, respectively.

## Author Contributions

Contribution: JB, BK, DZ, JDR, SSM, PV and LEP performed experiments. JB, BK, SSM and PV analyzed the data and prepared the figures. JB, BK, DZ, SSM, MK, BA and SKH were involved in designing the research and interpreting data. JB wrote the paper. BK, DZ, JDR, SSM, MK, BA and SKH edited the manuscript. All authors reviewed the paper.

## Code availability

Matlab based tissue kinetic fitting codes are available from the corresponding author upon reasonable request.

## Data availability

The data that support the findings of this study are available from the corresponding author upon reasonable request.

## Competing Interests

The authors declare no competing interests.

---

**Video 1:** Three-dimensional tile image of ALL bearing mouse

A MPM based three dimensional tiled image play-through of mouse calvarium 4 days post injection of GFP+ ALL. The z spacing between images is 15um. Bone-collagen (blue), ALL (green), and Blood pool (red) are shown.

**Video 2:** Time lapsed imaging of dextran kinetics

Time lapsed imaging of fluorescent TRITC dextran (gray) injection into irradiated ALL burdened mouse. Time post injection is displayed. Bone-Collagen signal is displayed to show bone marrow compartments within the calvarium (blue).

**Video 3:** Time lapsed imaging of Cellular clumping and intermittent BMV opening and closing

Time lapse dextran imaging (gray) of dextran injection into ALL (green) burdened mouse. Time post injection is displayed. Bone-Collagen signal is displayed to show bone marrow compartments within the calvarium (blue). Opening and closing of vessels and cell clumping is illustrated in the video.

**Video 4:** Time lapsed imaging of healthy control BMV

Time lapsed dextran imaging (gray) of dextran injection into healthy control mouse. Time post injection is displayed. Bone-collagen signal is displayed to show bone marrow compartments within the calvarium (blue). No opening or closing of vessels or cellular clumping is seen.

