## Supplemental Figures for "Development of Kinetic Modeling to Assess Multi-functional Vascular Response to Low Dose Radiation in Leukemia"

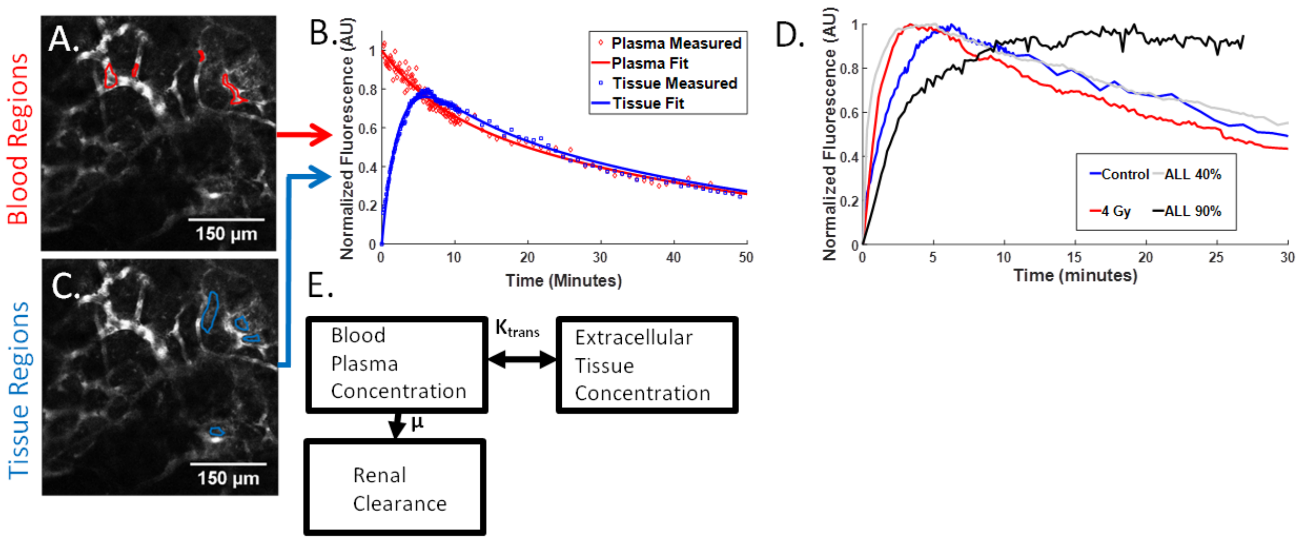


Supplemental Figure 1: Time lapsed dextran imaging and compartmental modeling.

TPM images showing contoured regions of interest for blood (red) and tissue (blue) mean fluorescent intensity quantification of dextran from a healthy control mouse (A,C). The corresponding blood and tissue data from images plotted with their fitting functions are shown (B). Normalized time-lapsed data for individual mice from control, moderate burden ALL, non-leukemic 4Gy, and high burden ALL mice (D). A diagram of the compartment model used for Dextran kinetics is shown (E).

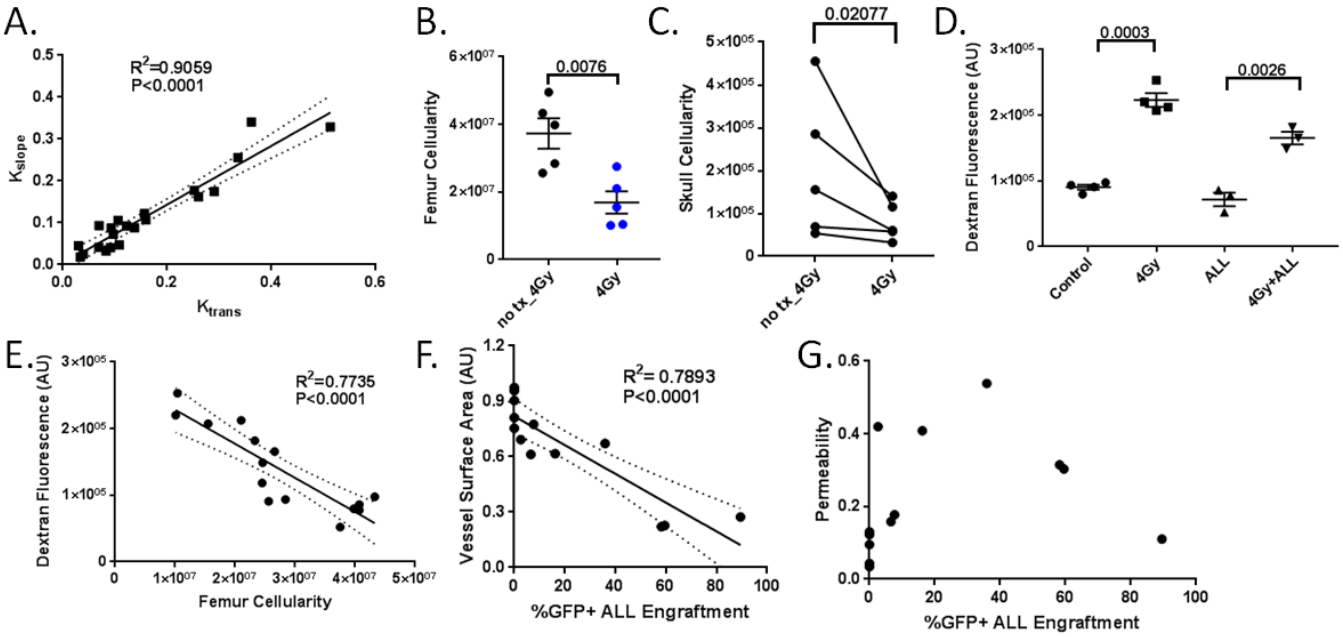


Supplemental Figure 2: Compartmental modeling validation and parameters.

A plot of $K_{trans}$ versus $K_{slope}$ from separate analysis methods of kinetic transfer coefficient are shown with R-squared value and best fit line (A). Total femur cellularity of treated and non treated femurs regions (B). Total skull cellularity in treated and non treated regions of healthy mice with a two sided paired t-test calculation (C.) Dextran fluorescence of crushed femur supernatants for healthy control, non leukemic 4Gy IGRT treated, ALL bearing, and 4Gy IGRT treated ALL bearing femurs (D). Femur cellularity versus dextran fluorescence of crushed femur supernatants with the corresponding R-squared value and best fit line (E). Vessel surface area versus percentage ALL engraftment with the corresponding R-squared value and best fit line are plotted for mice analyzed with tissue compartmental modeling (F). ALL bone marrow engraftment plotted versus vascular permeability for mice analyzed with tissue compartmental modeling (G).

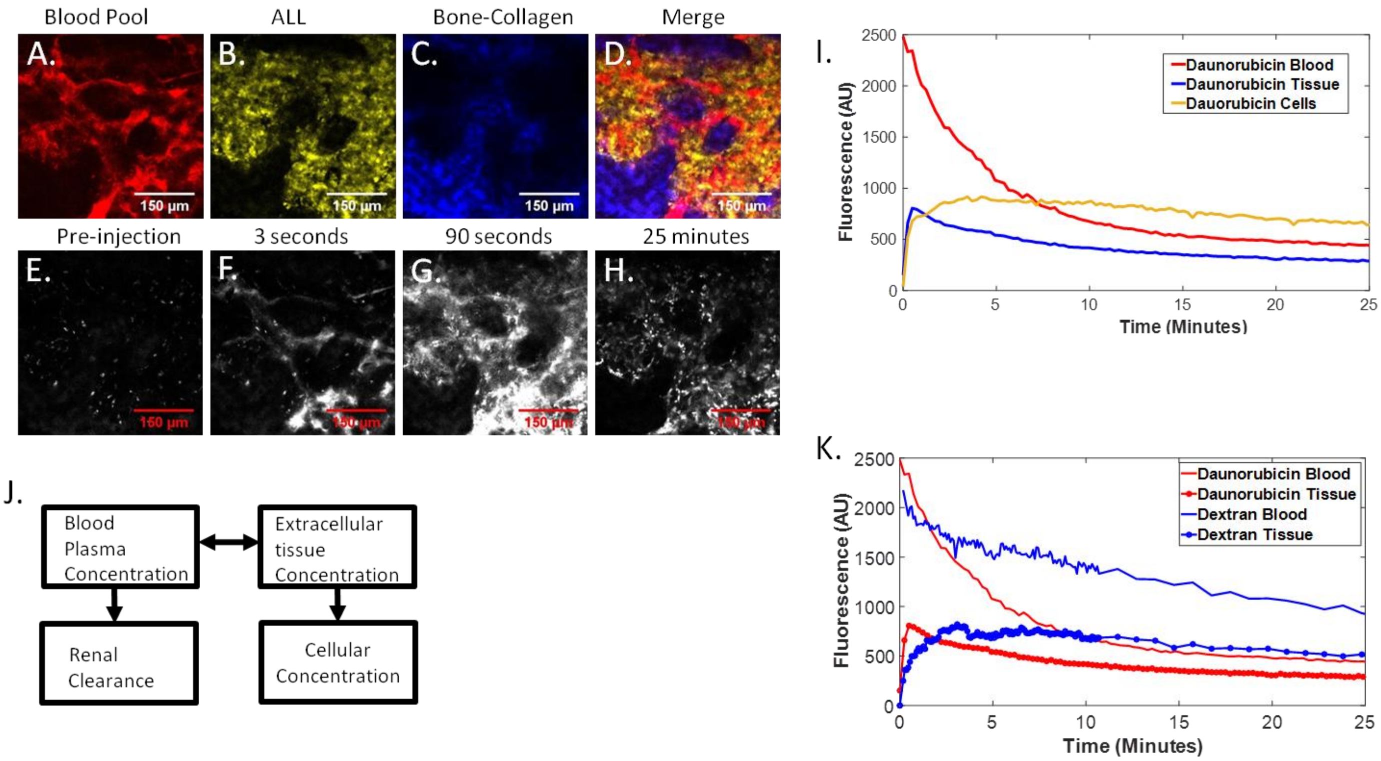


Supplemental Figure 3: Compartmental modeling of daunorubicin.

Corresponding ALL burdened mouse images of blood pool (A) ALL (B), bone-collagen(C), and a merged image (D) are shown for daunorubicin injected mouse. Images of daunorubicin fluorescence before (E), three second post (F) 90 second post (G) and 25 minute post (H) daunorubicin injection are shown. Clear distinction between blood, tissue and cellular compartments can be visualized. Plots of mean fluorescence versus time for contoured blood, tissue, and cellular compartments are shown (I). A representative image of a three tissue compartment model applicable to Daunorubicin is shown (J). A comparison between Daunorubicin and Dextran blood and tissue concentration curves for ALL burdened mice. (K).

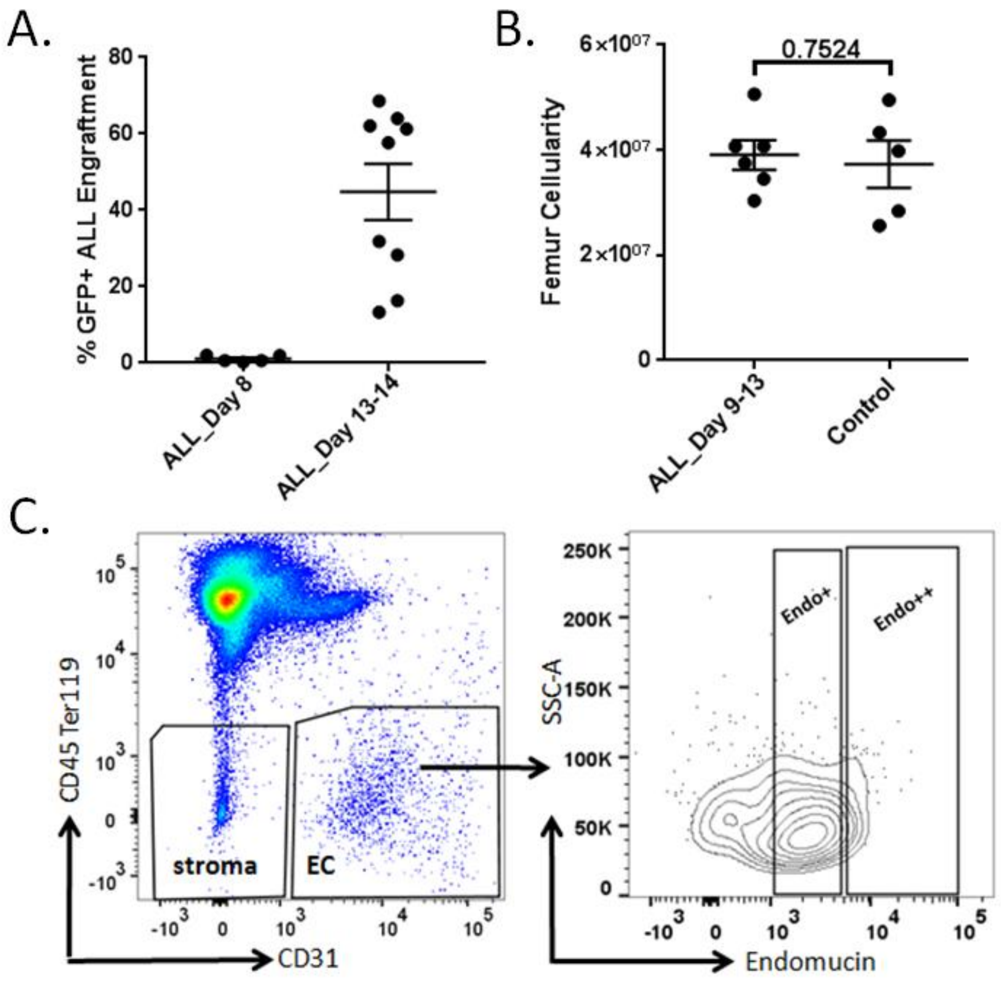


Supplemental Figure 4: Endothelial cell flow cytometry analysis.

Corresponding GFP+ ALL engraftment for mice analyzed for endothelial cell flow cytometry analysis in Figure 3F (A). Total cellularity of crushed femur bone marrow for healthy control mice and mice 9-13 days post injection of ALL (B). The flow cytometry gaiting scheme for CD31+ CD45Ter119- endothelial cell isolation and high endomucin subpopulation is shown (C).

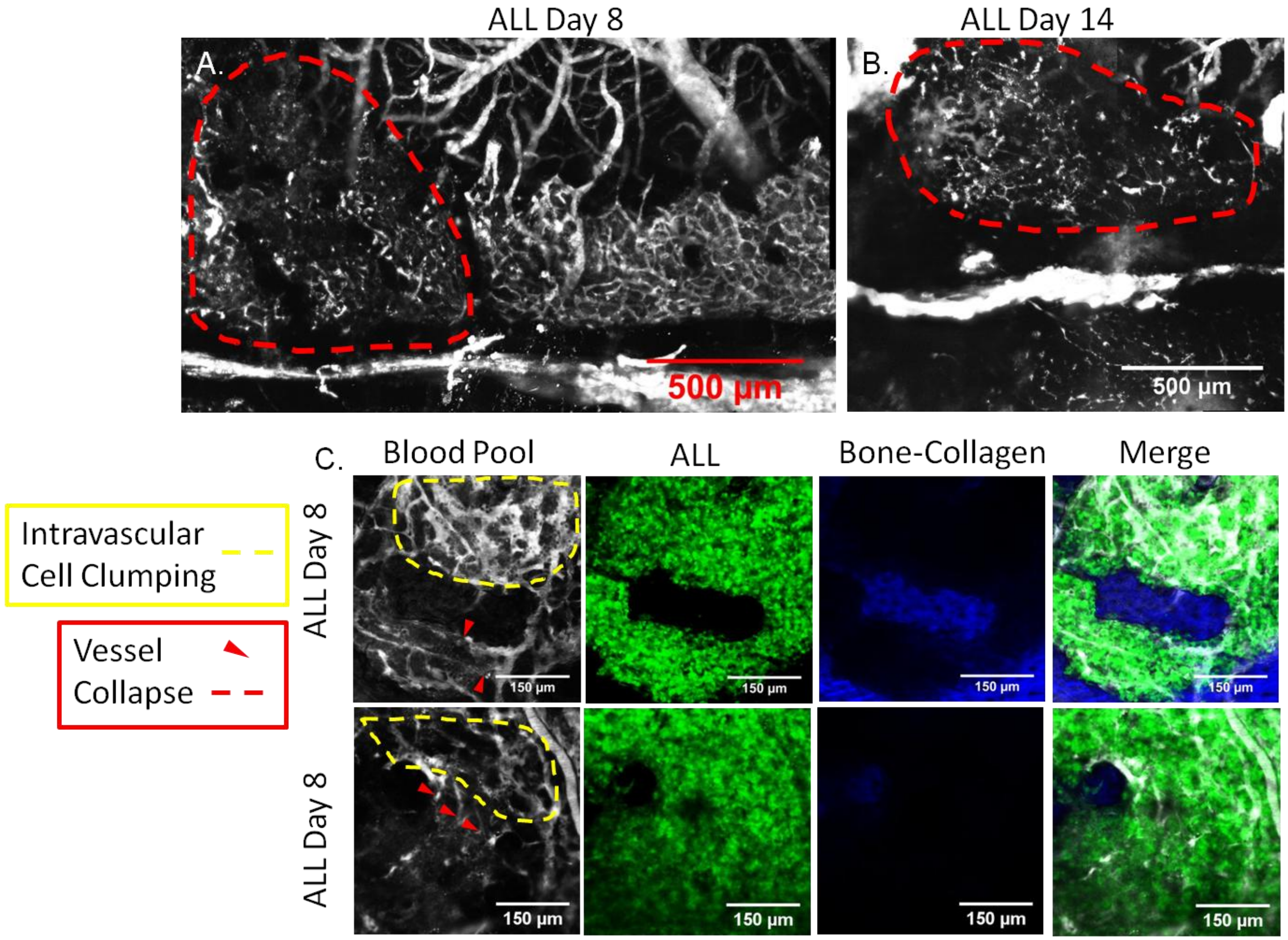


Supplemental Figure 5: Vascular collapse and intravascular cellular accumulation in ALL bearing mice.

MPM images of mouse calvarium vasculature blood pool in ALL transplanted mice 8 days (A) and 14 days (B) post ALL injection. Red dashed lines indicate regions of vascular collapse. Merged composite images of blood pool(gray), ALL (green), and collagen-bone(blue) for ALL injected mice 8 days post ALL injection(C). Red triangles and red dashed lines indicate specific vessel collapse and regions of vascular collapse respectively. Yellow dashed lines indicate regions of intravascular cell clumping.

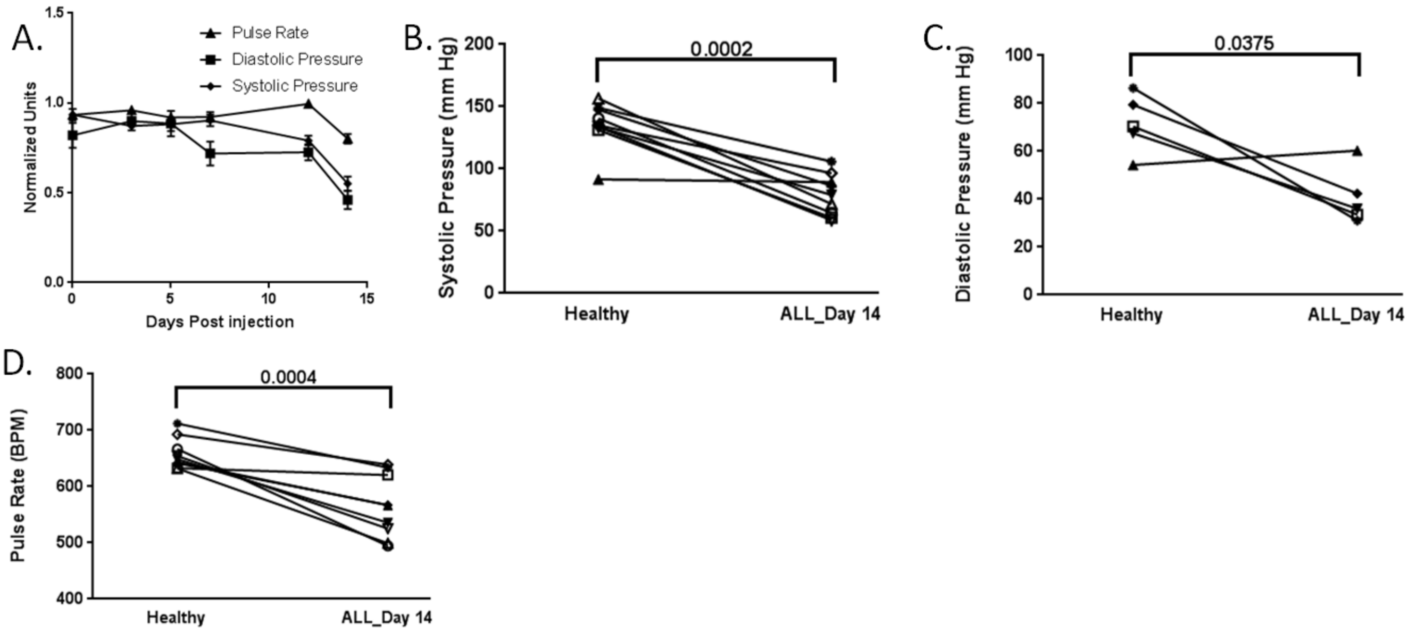


Supplemental Figure 6: Changes in systemic blood pressure with the onset of ALL.

A normalized plot of systemic mouse heart pulse rate, diastolic blood pressure, and systolic blood pressure at 3, 5, 7, 12, and 14 days following ALL injection (A). Plots of systolic blood pressure, diastolic blood pressure, and heart rate in mice before and 14 days after receiving ALL injection along with two sided paired T-test significance values (B-D).

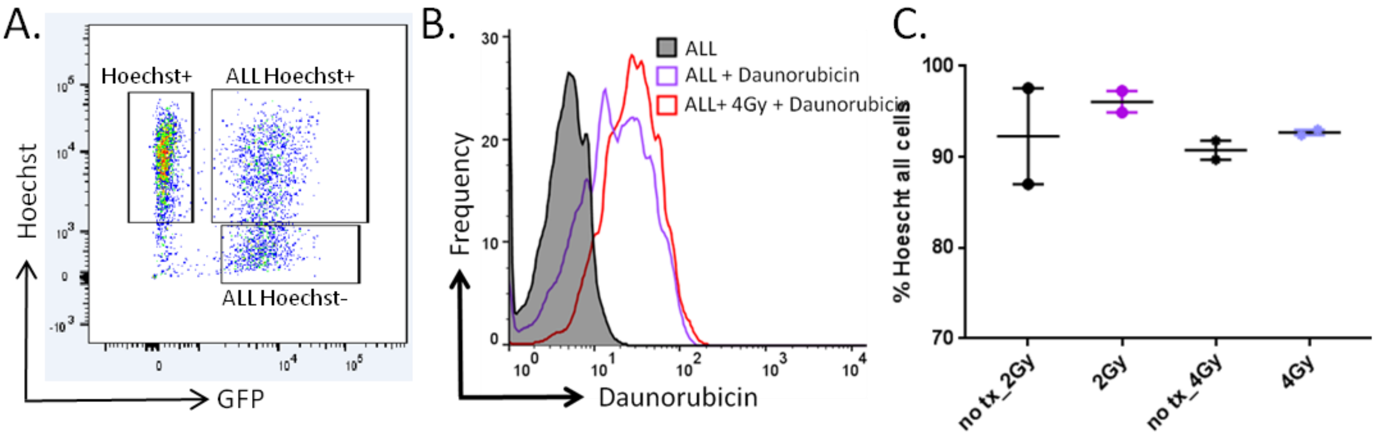


Supplemental Figure 7: Hoechst and daunorubicin uptake gaiting and analysis

A representative gating scheme showing clear separation between Hoechst positive, Hoechst positive ALL positive, and ALL positive Hoechst negative populations (A). A representative histogram of daunorubicin fluorescence in ALL cells for flow cytometry analysis is shown (B). The percentage of total skull cells stained positive for Hoechst in 2Gy IGRT and 4Gy IGRT healthy mice with their corresponding non treated regions (C).

Supplemental Video S1: Time lapsed imaging of ALL transendothelial migration

GFP+ Leukemia cell (green) homing out of blood pool (red) and into calvarium marrow space within 30 minutes of ALL injection is shown. A single ALL cell can be seen moving toward bone-collagen (blue).

Supplemental Video S2: Time lapsed imaging of daunorubicin

Time lapsed daunorubicin imaging (gray) of daunorubicin injection into ALL burdened mouse. Time post injection is displayed. Bone-collagen signal is displayed to show bone marrow compartments within the calvarium (blue).

Supplemental Video S3: Time lapsed imaging of interstitial cellular movement and BMV collapse

Time lapse dextran imaging (gray) of dextran injection into ALL (green) burdened mouse. Time post injection is displayed. Bone-Collagen signal is displayed to show bone marrow compartments within the calvarium (blue). Cellular movement and vessel collapse is illustrated in the video.
