## Supplemental Methods for "Development of Kinetic Modeling to Assess Multi-functional Vascular Response to Low Dose Radiation in Leukemia"

### *Mice*

8-10 week old female mice were used for endothelial cell analysis. Tile image acquisition and vessel morphology analysis were performed using 8-11 week old male and female mice. Vascular compartmental modeling, dextran well counter fluorescence, cellularity measurements, treatment engraftment measurements, and Hoechst dye flow cytometry were performed using 7-10 week old female mice. Intravascular cellular velocity measurements were performed using 7-12 week old female mice. Daunorubicin imaging and flow studies were performed with 10-12 week old male mice at moderate ALL burden (10-50%).

### *Multiphoton Image Acquisition and Image Processing*

An Olympus XLUMPlanFL 20x objective (1.00 NA water objective) was used for image acquisition of all images. Simultaneous four channel acquisition was performed at 660/40 (far red), 595/50 (red), 525/50 (green), and 460/50(blue) for visualization of leukemia, and blood pool agents. Data acquisition using the Prarie Ultima microscope was handled by Prairieview 5.3 software. Imaging excitation was performed at 900nm for $GFP^+$ ALL, TritC Dextran, and Qtracker™ 655 Vascular Label. Additional excitation was performed at 840nm and 960 nm for visualization of both Daunorubicin and $GFP^+$ ALL respectively. Color channel windows were adjusted individually in Fiji/ImageJ to improve image display quality. Individual color window adjustments were identical between sets of comparative images.

20µl of Qtracker™ 655 Vascular Label was added to 180µl of PBS and administered through tail vein injection approximately 10 minutes before imaging as a blood pool imaging agent for tiled image sets, vascular diameter measurements, and vascular density measurements. Tiled images were acquired by collecting approximately a 5x4 grid series of overlapping z-stack images. The images were overlapping by 15% with a resolution of 512 by 512 pixels and a z-slice spacing of 15um. Tiled images were stitched together using a Fiji/ImageJ grid collection/stitching plugin[1]. Vascular diameter and the number of vessel branches per area were quantified using Fiji/ImageJ.

Image subtraction and cellular segmentation was performed using Cell Profiler. Image subtraction was used for quantitative ALL and Daunorubicin fluorescence analysis as well as all images presentation for all images except time lapsed Daunorubicin image sets. For 900nm excitation ALL image display and quantification, auto fluorescence from blue and red channels were multiplied by a scalar and subtracted from the green channel to eliminate signal from both collagen and auto fluorescent cells. For Daunorubicin based single time-point images, image subtraction was performed by multiplying the 840nm excitation blue channel signal by a scalar and subtracting from the 960nm excitation green channel and 840nm excitation red channel to identify $GFP^+$ ALL and Daunorubicin signal respectively. Healthy mice without Daunorubicin injections were used to determine image subtraction scalars.

### *Intravascular Cellular Velocity Measurements*

Vessel cellular velocities were measured after injection of either 150 kDa TRITC Dextran or Qtracker 655 Vascular labels. Cells in the bloodstream could be visualized as shadows passing through the blood plasma. Repeated line scans were taken on the center of blood vessels with frequencies of approximately 0.4 to 3 kHz. Plots of the time-space images were acquired where cells moving through the vessels are plotted as diagonal lines. The average slope of 3 separate lines was found using Fiji/ImageJ to calculate cellular velocity.

### *Vascular Permeability and Compartmental Modeling*

Immediately following injection, time lapsed single or z-stack images of Dextran leakage into the tissue were taken initially with 6-15 second intervals, increased to 60 seconds after peak concentration had been reached. Blood and tissue regions of interest were contoured in Fiji/ImageJ by attempting to contour regions of maximized total signal for both blood and tissue. When needed, contours were shifted frame by frame to account for small amounts of lateral drift. Dextran fluorescent intensity was measured after spectral unmixing of GFP+ BCR-ABL cellular bleed through into Dextran signal calculated using equation 1.

$$\mathrm{I_D}(\mathrm{t}) = \frac{\mathrm{R(t)} - \mathrm{R_B} * \frac{\mathrm{G(t)}}{\mathrm{G_B}}}{\mathrm{A}} \qquad [1]$$

Where R(t) and G(t) are the total red and green channel signal intensity at time t for the region of interest. $R_B$ and $G_B$ are the background signal intensities prior to dextran injection for the red and green channels respectively. A is the area of the region of interest being analyzed. The equation for spectral unmixing was used to account for photobleaching of GFP+ ALL cells during time lapsed imaging. Daunorubicin time lapsed quantification was performed using pre-injected signal background subtraction rather than spectral unmixing as small amounts of Daunorubicin signal was present in the green channel.

Dextran transport from tissue and blood regions of interest was modeled using Equation 2.

$$\frac{dC_t}{dt} = (K_{trans})(C_p(t) - v_{ec} * C_t(t)) \qquad [1]$$

Where $C_p(t)$ is the capillary plasma tracer concentration, and $C_t$ is the tracer tissue concentration. A value of 0.4 for the hematocrit was used to convert blood concentration to $C_p(t)$. $C_p(t)$ was modeled by a summation of two separately weighted exponential decay curves and fit to each individual animals blood plasma signal. Curve fitting was performed in Matlab using the lsqnonlin() function. Equation 2 was solved for by substituting the blood plasma equation and setting the initial tissue concentration equal to zero.

$$C_t(t) = K_{trans} \sum_{i=1}^{n} \left[ \left( \frac{A_i}{(K_{trans}/\nu_{ec}) - \mu_i} \right) \left( e^{-\mu_i * t} - e^{-(K_{trans}/\nu_{ec}) * t} \right) \right] \qquad [3]$$

Where $A_i$ and $\mu_i$ are the scaling coefficient and attenuation coefficient for the i$^{th}$ exponential plasma concentration curve respectively. $A_i$ and $\mu_i$ are used to fit equation 3 to tissue data, solving for $K_{trans}$ and $v_{ec}$. The fitting was performed from a range of starting conditions across the acceptable boundary conditions to ensure the error function was globally minimized.

$K_{trans}$ can additionally be measured by observing the rate of change of $C_t$ immediately following an injection of a bolus before any dextran backflow from the tissue to the vessels can occur. [2-4] Immediately following injection $C_t(0) \approx 0$ which, when substituted into equation 2, gives

$$K_{slope} = \frac{\frac{dC_t}{dt}}{C_p} = \frac{Slope_{initial}}{C_p} \qquad [4]$$

*Slope*$_{initial}$ was calculated by taking the difference in mean signal intensity before and after the first 18-30 second increment of data that showed a semi-linear increase in tissue concentration. This was divided by the time that elapsed between measurements. The blood plasma concentration during the slope was found by averaging four frames after the initial first pass of dextran marrow had equilibrated. Since calculations of $K_{slope}$ do not allow solutions for $v_{ec}$, primary analysis was performed using the curve fitting methodology obtaining $K_{trans}$ and $v_{ec}$.

It can be shown $K_{trans}$ is approximately equal to the average vascular permeability times the average vascular surface area per volume of tissue when the majority of first pass Dextran through the BMV remains in the vasculature and does not entirely permeate into the tissue bed [4]. Vascular permeability was calculated by dividing $K_{trans}$ by vessel surface area per imaging volume. The length and diameter of the lumen for each vessel was measured manually in Fiji/ImageJ and the respective surface area was calculated by approximating vessels as perfect cylinders. The amount of available calvarium marrow space in each imaging area was accounted for by observation of bone-collagen SHG to visualize the calvarium marrow space.

### *Blood Pressure Measurements*

Mouse blood pressure was recorded in mice using the BP-2000 Blood Pressure Analysis System (Vistech Systems). 30 successive tail cuff measurements were taken on each day. The first 10 measurements were discarded to allow time for mice to grow accustomed to the tail cuff. Successful measurements from the last 20 measurements were averaged together to obtain results.

### *CT Image Acquisition and Targeted Irradiation*

CT skull imaging was performed with 40kVp, 1mA, and a 100um resolution. The edge of an anterior 10mm square beam treated the top of the skull with a soft tissue equivalent dose of 2Gy or 4Gy (approximately 5Gy or 10Gy to bone respectively). The beam was placed 200um to the mouse's right side from the sagittal suture and completely covered the parietal bone on the mouse's left side. The left

Femur was treated with a 10mm square anterior to posterior and posterior to anterior beam (AP:PA). Dose was normalized to 2Gy or 4Gy 2mm below the surface of the skull for parietal beam and 2Gy or 4Gy within the marrow for the femur beam. Treatment was performed at 255kVp and 13mA.

### *Flow Cytometry*

For bone marrow endothelial cell analysis, the bones (Tibias and Femur) from normal and ALL mice were crushed, incubated with ACK lysis buffer(Gibco) for 5-10 min and washed with PBS. The bone fragments were incubated with collagenase (3mg/ml, Sigma Aldrich St Louis, MO) for 45min at 37°C to liberate the leukemic and bone marrow microenvironment cells from the bone surface [5]. The cells were filtered through a 40 µm filter and centrifuged at 300xg for 5min with PBS (.1%BSA) solution at 4°C. The cells were then stained and washed in PBS (.1% BSA). Stained cells were analyzed on FACSARIAIII (Bdbioscience, Franklin Lanes, NJ) and analyzed in live non-leukemic (GFP+) CD45-Ter119-CD31+ fraction. The flow analysis was performed using FlowJo 10.5.3 software (FlowJo, Ashland, OR).

For Hoechst uptake, treated and untreated calvarium samples were crushed and incubated with ACK lysis buffer to remove RBC. The PBS suspended cells were analyzed for Hoechst uptake in both the total cell and GFP+ ALL cell populations using the BD Fortessa (Bdbioscience, Franklin Lanes, NJ) on the same day.

For Daunorubicin uptake, treated and untreated femur samples were crushed and incubated with ACK lysis buffer to remove RBC. The cells were stained for viability amcyan dye (1µg/ml) for 15min and washed with PBS. Cells were analyzed on Amycan-(live) GFP+ leukemic cells population using BD FACSARIAIII (Bdbioscience, Franklin Lanes, NJ) sorter 2-3 hours post-harvest and were kept on ice throughout the staining process.

### *Dextran Well Plate Readings*

Dextran femur fluorescence was measured in imaged anesthetized mice euthanized 50 minutes post injection. Femurs were crushed in 7ml of PBS and centrifuged at 1200 RPM for 5 minutes. 200µl of supernatant was removed and read using a FilterMax F5 Multi-Mode Microplate Reader (Molecular Devices San Jose, CA). Well plate readings were linear over the read range (data not shown).
